# A Toxin-Antidote Selfish Element Increases Fitness of its Host

**DOI:** 10.1101/2022.07.15.500229

**Authors:** Lijiang Long, Wen Xu, Annalise B. Paaby, Patrick T. McGrath

## Abstract

Selfish genetic elements can promote their transmission at the expense of individual survival, creating conflict between the element and the rest of the genome. Recently, a large number of toxin-antidote (TA) post-segregation distorters have been identified in non-obligate outcrossing nematodes. Their origin and the evolutionary forces that keep them at intermediate population frequencies are poorly understood. Here, we study a TA element in C. elegans called peel-1/zeel-1. Two major haplotypes of this locus, with and without the selfish element, segregate in C. elegans. Here we study the fitness consequences of the peel-1/zeel-1 element outside of its role in gene drive in non-outcrossing animals. We demonstrate that loss of the toxin peel-1 decreased fitness of hermaphrodites and resulted in reductions in fecundity and body size. This fitness advantage is independent of the antidote zeel-1, suggesting that a distinct peel-1 pathway plays a biological role. This work demonstrates that a TA element can provide a fitness benefit to its hosts, either during their initial evolution or by being co-opted by the animals following their selfish spread. These findings guide our understanding on how TA elements can remain in a population where gene drive is minimized, helping resolve the mystery of prevalent TA elements in selfing animals.

## INTRODUCTION

Selfish genetic elements, or selfish genes, are heritable segments of DNA that promote their own transmission relative to the rest of the genome, potentially at the expense of the individual organism (Werren, 2011; Werren et al., 1988). They act through a diverse catalog of molecular mechanisms to increase their frequency, including transposons, homing endonucleases, sex-ratio distorters, and segregation or post-segregation distorters (Hurst & Werren, 2001). Because selfish genetic elements induce tension between genes and the hosts that carry them, including causing disease and other health problems, their discovery and study over the last 50 or so years has motivated major questions—and debate—over the nature and consequences of genetic conflict in inheritance systems (Ågren, 2016; Ågren & Clark, 2018; Hurst & Werren, 2001). In an early review, and in its revisit 23 years later, Werren and colleagues (2011; 1988) posed three questions about selfish genetic elements that remain outstanding today: (i) how they arise, (ii) how they are maintained, and (iii) how they influence evolution.

Theory and observation have indicated that selfish genetic elements decrease in prevalence as inbreeding in a system increases; spreading necessarily requires outcrossing to a vulnerable genetic background (Ågren & Clark, 2018; Hurst & Werren, 2001). However, a recent wave of discovery of toxin-antidote (TA) elements in non-obligate outcrossing species (e.g. Ben-David et al., 2017, 2021; Noble et al., 2021; Nuckolls et al., 2017; Shen et al., 2017) challenges this view. TA elements are post-segregation distorters composed of two or more linked sub-elements, including a “toxin” transmitted cytoplasmically from the parent to the offspring through the gamete and an “antidote” that rescues when expressed in the zygote. TA elements induce heavy fitness costs to hybrids heterozygous for an active/inactive genotype because while all gametes will carry the cytoplasmic toxin, only those zygotes that inherit the TA allele will express the antidote and survive.

TA systems, also referred to as “gamete killers” (e.g. Nuckolls et al., 2017) or Medea elements (e.g. Beeman et al., 1992; Noble et al., 2021), have been identified across multiple kingdoms of life, including bacteria, plants, fungi, insects, and nematodes (Akarsu et al., 2019; Bardaji et al., 2019; Beckmann et al., 2017; Beeman et al., 1992; Ben-David et al., 2021; Chen et al., 2008; Leplae et al., 2011; Saavedra De Bast et al., 2008; Seidel et al., 2011; Yang et al., 2012). In the nematode genus *Caenorhabditis*, androdioecy (male and hermaphrodite sexes) has evolved independently three times from a male-female ancestor (Ellis, 2017); consequently *C. elegans, C. briggsae* and *C. tropicalis* reproduce primarily by selfing, with infrequent instances of outcrossing via male mating (Barrière & Félix, 2005; Cutter et al., 2006; Noble et al., 2021). TA elements have been identified in all three species, including multiple elements in both *C. elegans* and *C. tropicalis* (Ben-David et al., 2017, 2021; Noble et al., 2021; Seidel et al., 2008, 2011). Similar elements have not been identified in obligate outcrossing *Caenorhabditis* nematodes. These results beg the question: Why have so many TA elements been identified in non-obligate outcrossing species (Noble et al., 2021; Sweigart et al., 2019)?

One of the most complete mechanistic descriptions of a TA system is the *zeel-1;peel-1* locus in *C. elegans*, in which a sperm-delivered toxin (*peel-1*) induces arrest in embryos not carrying the zygotically expressed antidote (*zeel-1*) (**Figure 1A**) (Seidel et al., 2008, 2011). The alternative active/inactive haplotypes that segregate within *C. elegans* exhibit high genetic diversity (**Figure 1B**) that dates the divergence of the two haplotypes to roughly 8 million generations ago (Seidel et al., 2008). Maintenance (**Figure 1C**) of ancient polymorphism is inconsistent with a history of selfish activity: in outcrossing populations, genic drive should fix the active haplotype rapidly; in the androdioecious mating system of *C. elegans*, a high rate of selfing should fix an element at high frequency or allow it to be lost by drift at low frequency (Noble et al., 2021). However, it is unknown how the fitness of a TA element, independent of its selfishness, may influence its spread or maintenance.

**Figure 1.**
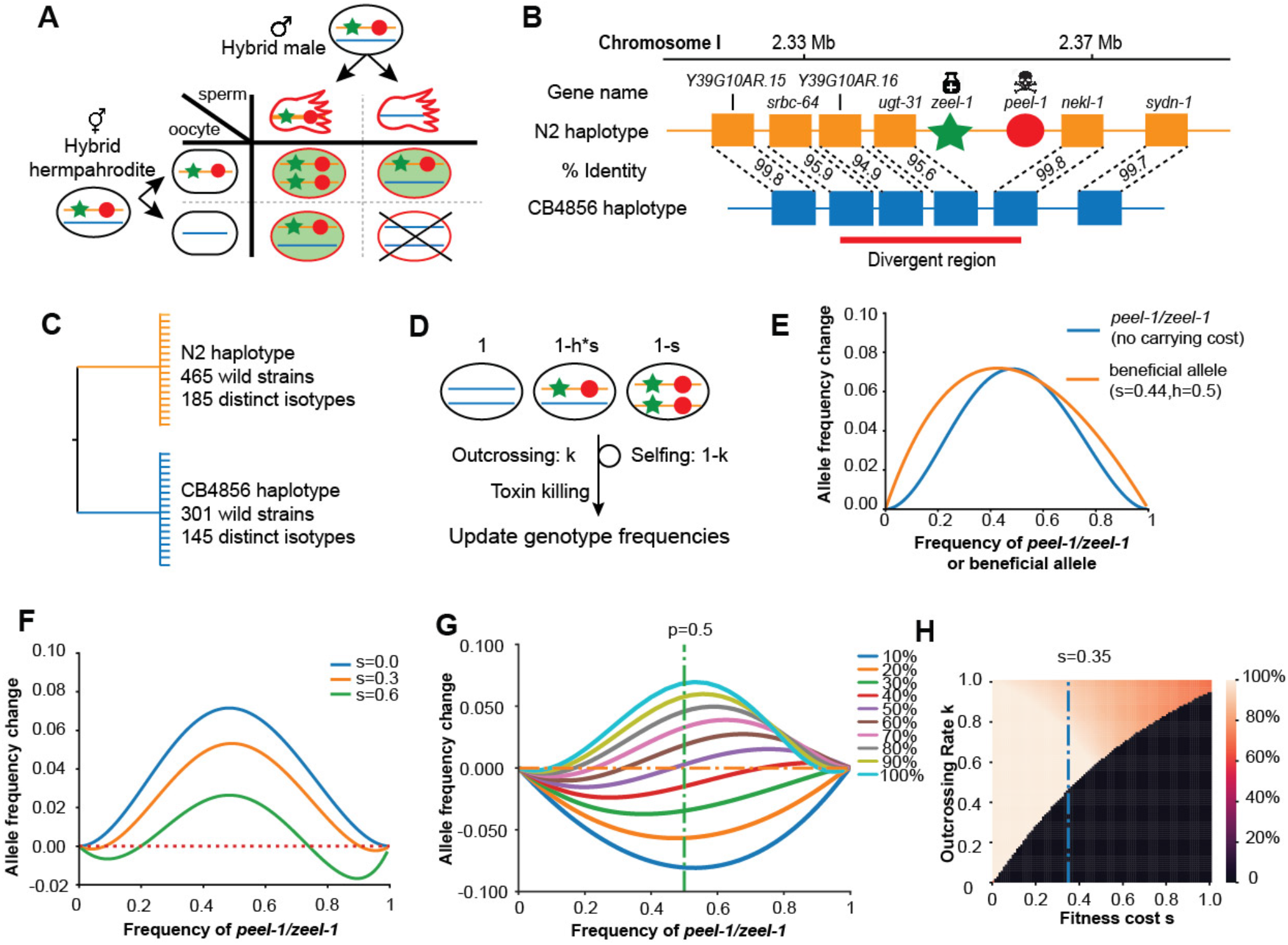
Description and models of selection for *peel-1/zeel-1.* **A.** Schematic of the progenies created from an F1 hybrid cross, produced through intercrossing. Red outline indicates cytoplasmic inheritance of the PEEL-1 toxin from the hybrid male, independent of genomic inheritance of *peel-1* (red circle) or *zeel-1* (green star), which counteracts the toxin by zygotic expression (green background). Progeny that die are indicated by the X cross. **B.** Schematic of the genomic region surrounding *zeel-1;peel-1* for two major haplotypes, N2 and CB4856. *peel-1/zeel-1* is present in the N2 genome and deleted in the CB4856 genome. Amino acid identities of each gene are shown between the two haplotypes. The red bar denotes the hyperdivergent region starting in the 5’ end of *srbc-64* and ending in the beginning of *nekl-1.* **C.** A gene tree representation of the *peel-1/zeel-1* locus from wild strains of *C. elegans* using the hyperdivergent region (based on Seidel et al., 2008). Two major branches distinguish the N2 and CB4856 haplotypes; the number of wild isolates and distinct isotypes are labeled on each branch. This distribution is consistent with balancing selection acting on each haplotype. **D.** Schematic of the simulation of *peel-1/zeel-1* population dynamics. The fitness of each genotype is shown on top. Genotype frequencies are updated each generation using Table S1. **E.** The allele frequency change per generation (y-axis) of *peel-1/zeel-1* (s=0, k=1, blue curve) or a beneficial allele (s=0.44, h=0.5) as a function of allele frequency (x-axis). **F.** The change in allele frequency per generation (y-axis) of *peel-1/zeel-1* with three different carrying costs (s=0, s=0.3, and s=0.6), as a function of allele frequency (x-axis). **G.** The change in allele frequency per generation (y-axis) of *peel-1/zeel-1* with a fixed fitness cost (s=0.35, h=0.5) at different rates of outcrossing, as a function of allele frequency (x-axis). **H.** Heat map showing the *peel-1/zeel-1* frequency after 1000 generations, over varying outcrossing rates (y-axis) and carrying costs (x-axis). Initial frequency of the element was 50%. Black indicates animals that have lost the element.

In this study, we investigate the fitness effect of a TA element in the host genotype, independent of its toxic incompatibility in outcrossed individuals, to assess its role in maintaining the prevalence of TA elements in non-obligate outcrossing populations. Modeling under expected conditions shows that TA elements are vulnerable to being lost at low frequency, but direct tests of fitness-proximal traits indicate that the active *peel-1* allele increases fitness relative to the inactive haplotype. These results suggest that the spread of the *zeel-1;peel-1* allele within *C. elegans* might not be gene drive, but positive selection acting on independent biological traits. These findings have consequences for considering the origin and maintenance of TA elements and their influence on the historical evolution of populations.

## RESULTS AND DISCUSSION

### The fitness cost of a TA element influences its initial spread and final fate

The effectiveness of a gene drive system is dependent on multiple factors beyond its selfish induction of incompatibility, including genotype frequency, outcrossing rate, and fitness in the host background. To explore these parameters, we adapted a family-based model (**Figure 1D, Table S1**) (Wade & Beeman, 1994) with modifications to account for paternal delivery of the toxin, the androdioecious mating system of *C. elegans*, and selection cost of the element.

Under a simple scenario of no fitness consequence to the host genotype (s=0) and a completely outcrossing population (k=1), the element spreads rapidly through the population with a maximum allele change comparable to an additive beneficial allele with a selection coefficient of 0.44 (**Figure 1E**), two to four times higher than the selection coefficient of lactase persistence in humans (Bersaglieri et al., 2004). However, gene drive is weaker than the beneficial allele at the tails of the allele frequency range: at low frequency, the rarity of the element limits how fast it spreads; at high frequency, the rarity of the vulnerable genotype slows its approach to fixation. If the element induces a carrying cost to the host genotype (e.g., s=0.3, s=0.6), for example via energy expenditure or “leaky” toxicity, the dynamics at the extreme allele frequencies are amplified (**Figure 1F**). At low frequency, the carrying cost counteracts gene drive, reducing the likelihood that the element reaches appreciable frequency by genetic drift before being lost. At high frequency, the carrying cost compounds the slowing rate of gene drive such that it reaches a stable equilibrium and does not fix.

Previous models have shown that spread of a TA element accelerates with the rate of outcrossing (Noble et al., 2021). Given a substantial carrying cost to the host genotype (s=0.35), a TA element is likely to increase in frequency only under relatively high rates of outcrossing (**Figure 1G**). Under outcrossing rates typical for *C. elegans* (Barrière & Félix, 2005; Sivasundar & Hey, 2005), the element will likely to be lost from the population under all but the mildest carrying costs (**Figure S1**), as increasing fitness costs require increasing outcrossing for the element to reach a stable equilibrium (**Figure 1H**).

Given these dynamics, we are challenged to explain how a novel TA element could rise in initial frequency in a population. One hypothesis is that TA elements in non-obligate outcrossing *Caenorhabditis* may have originated in an outcrossing ancestor, then persisted by other evolutionary forces such as drift or balancing selection (Noble et al., 2021; Seidel et al., 2011; Sweigart et al., 2019). Such a scenario is consistent with the recent opinion by Sweigart and colleagues (2019), who argue that TA elements may exist in nature with only incidental instances of “selfish” activity. This shift away from the conventional framing of TA elements as consistently selfish makes sense in the context of non-obligate outcrossing populations, which permit elements to proliferate in sequestered lineages without conflict.

### The active zeel-1;peel-1 haplotype is associated with higher fitness in laboratory environments

To investigate its potential to spread through the population without conflict, we evaluated the fitness consequences of the *peel-1/zeel-1* element independent of its incompatibility cost in heterozygotes. First we employed a previously described fitness assay (Large et al., 2016; Zhao et al., 2018) to compete N2^*zeel-1;peel-1(CB4856)*^, which carries a ~140-370kb interval spanning the *zeel-1;peel-1* locus from CB4856 introgressed into N2 (Ben-David et al., 2017), against N2^*marker*^, a modified version of N2 carrying a silent marker mutation in the *dpy-10* gene. As CB4856 harbors the inactive haplotype, N2^*zeel-1;peel-1(CB4856)*^ lacks the toxin/antidote element, while N2^*marker*^ carries the active element native to N2. In these assays, males are not present and outcrossing is prevented, so relative fitness is estimated from true-breeding hermaphrodite genotypes.

N2^*marker*^ outcompeted N2^*zeel-1;peel-1(CB4856)*^ (**Figure 2A**), with a relative fitness (w) of 1.18 (1.15-1.21, 95% CI). Association of the active allele with higher fitness suggests that induction of *peel-1* toxicity and/or rescue by *zeel-1* is not costly, that the active allele is linked to one or more mutations in the N2 background that confer an independent fitness advantage, or both. These mutations could reside within *zeel-1;peel-1*, within the four nearby genes within the high diversity region, or outside the high diversity region but within the 140-370kb introgressed region of this strain (**Figure 1A**). We also measured fecundity and body size in N2 and N2^*zeel-1;peel-1(CB4856)*^ directly, and observed similar outcomes: N2 laid 9% more embryos (p<0.001, **Figure 2B**) and was 9% larger 72 hours after hatching (p<0.001, **Figure 2C**), indicating a faster growth rate.

**Figure 2.**
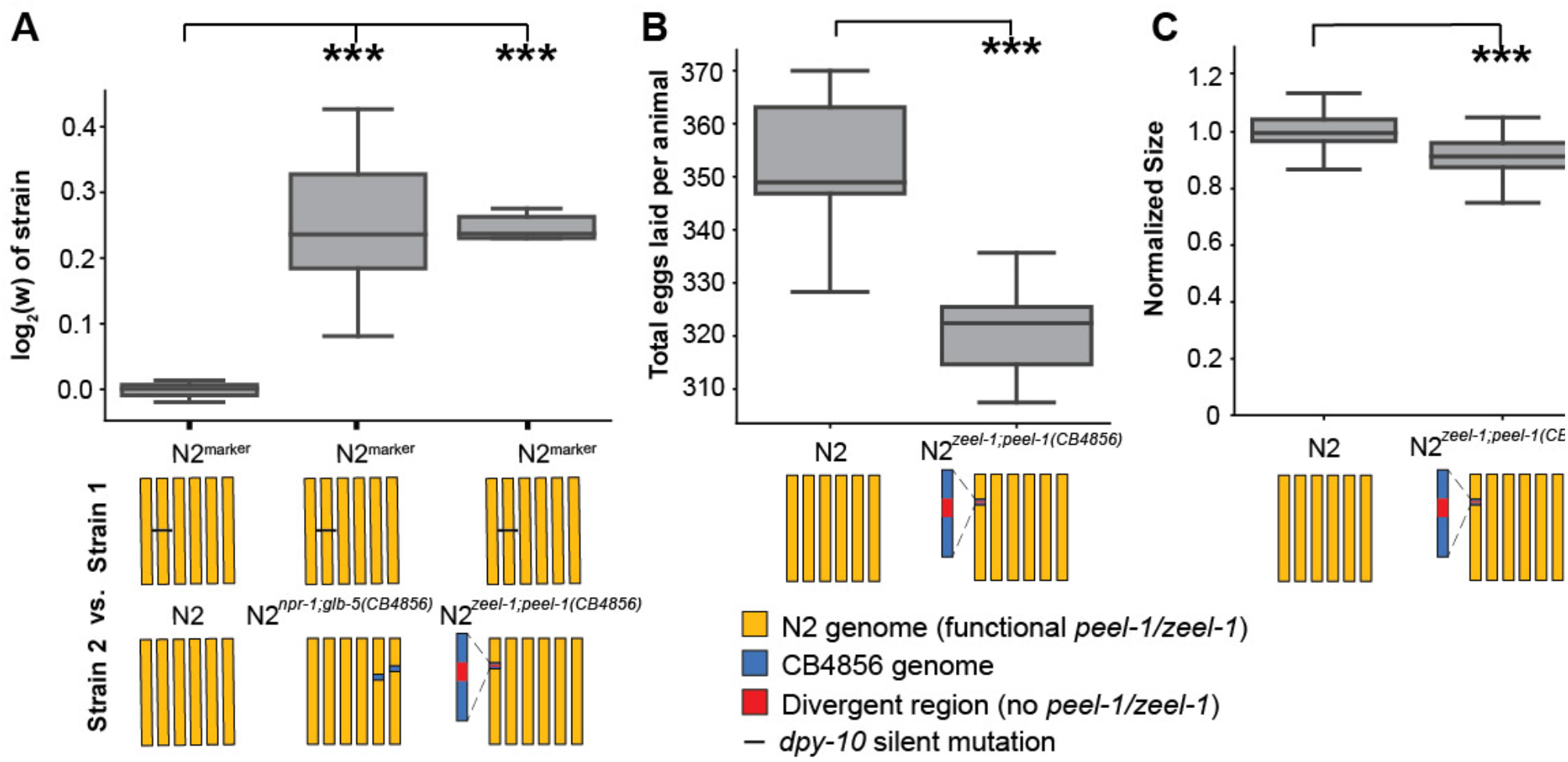
*peel-1/zeel-1* is linked to genetic variation that increases fitness in the host genotype in laboratory conditions. **A.** Relative fitness of experimental genotypes competed against N2^*marker*^, which has a silent mutation in *dpy-10* used as a barcode for digital PCR. N2^*marker*^, which has the *peel-1/zeel-1* element native to N2, outcompeted N2^*zeel-1;peel-1(CB4856)*^, which has a ~140-370kb interval spanning the *zeel-1;peel-1* locus from CB4856 introgressed into N2 (Ben-David et al., 2017). The relative fitness of N2^*marker*^ over N2^*zeel-1;peel-1(CB4856)*^ (w = 1.18, 1.15-1.21, 95% CI) is similar to its relative fitness over N2^*npr-1;glb-5(CB4856)*^ (w = 1.19, 1.10-1.28, 95% CI), which was used as a positive control. N2^*npr-1;glb-5(CB4856)*^ carries introgressed CB4856 alleles at *npr-1* and *glb-5* that were previously shown to decrease fitness relative to N2 alleles in laboratory conditions (McGrath et al., 2009). The relative fitness of N2 versus N2^*marker*^ is not significantly different than zero, indicating that the *dpy-10* barcode allele in N2^*marker*^ does not affect fitness. **B.** Fecundity of N2 and N2^*zeel-1;peel-1(CB4856)*^. **C.** Growth/size analysis of N2 and N2^*zeel-1;peel-1(CB4856)*^. The body size of young adult animals were measured at 72 hours and normalized to the average size of N2. For all plots, the box plot shows quartiles of the dataset while the whiskers cover the entire distribution of the data minus outliers. ***p<0.001 by two-tailed t-test.

These results indicate that variants associated with the active *zeel-1;peel-1* haplotype promote fitness in the host genotype, providing a potential mechanism for proliferation and persistence of the element in selfing lineages.

### The active peel-1 allele is associated with higher fitness in laboratory environments

To test the fitness consequences of the *peel-1* toxin directly, we used CRISPR/Cas9 to engineer a knock out of *peel-1* in the N2 background. N2^*peel-1(null)*^ produces a truncated protein of 46 amino acids (relative to 174) via an early stop codon (**Figure 3A**). We verified loss of function by embryo killing assays: N2 crossed to CB4856 produced the expected 25% embryonic lethality from selfed F1 hermaphrodites; the N2^*peel-1(null)*^ cross produced zero dead embryos (**Figure 3B**). Interestingly, the *peel-1(null*) allele affected fitness proximal traits and fitness in laboratory conditions. The N2^*peel-1(null)*^ produced 6% fewer offspring (**Figure 3C**) and were 7% smaller 72 hours after hatching than N2 (**Figure 3D**). Competition experiments between N2^*peel-1(null)*^ against N2^*marker*^ or N2^*peel-1(null),marker*^ against N2 also demonstrated a fitness increase associated with the active *peel-1* allele (w = 1.06, 1.04-1.07, 95% CI) (**Figure 3E**); this fitness difference accounts for 32% of the difference arising from the N2^*zeel-1;peel-1(CB4856)*^ comparison. Thus, while *peel-1* acts as a toxin in the context of outcrossing cross-progeny, it increases the fitness of selfing hermaphrodites in laboratory conditions. These results suggest that *peel-1* is not simply a toxin gene, and plays some other biologically relevant role in *C. elegans.*

**Figure 3.**
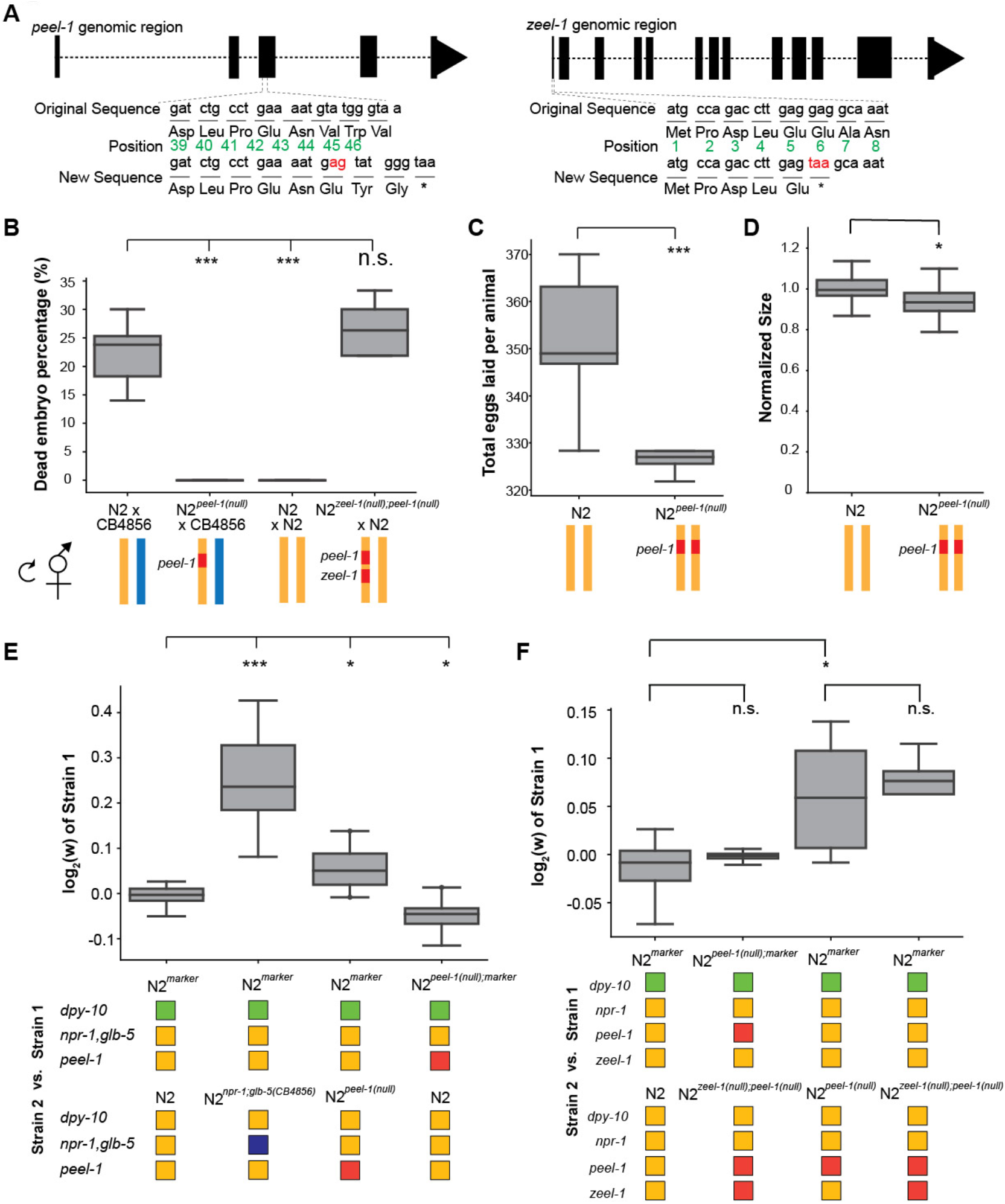
Tests of *peel-1* and *zeel-1* function using CRISPR/Cas9 show *peel-1* increases fitness independent of *zeel-1.* **A.** Schematic of the *peel-1* and *zeel-1* loss-of-function alleles. At *peel-1*, two additional nucleotides (marked in red) inserted into the third exon generate a frameshift and an early stop codon (marked by *). The green numbers denote the amino acid position of the PEEL-1 protein sequence. For *zeel-1*, a two-nucleotide replacement induces an early stop codon (marked by *). **B.** N2^*peel-1(null)*^ and N2^*zeel-1(null);peel-1(null)*^ carry loss-of-function alleles, as selfed cross-progeny show: N2 x CB4856 produce ~25% embryonic lethality, but N2^*peel-1(null)*^ x CB4856 produce 0%; N2 x N2 produce 0%, but N2^*zeel-1(null);peel-1(null)*^ x N2 produce ~25%. **C.** Fecundity of the N2 and N2^*peel-1(null)*^ strains. **D.** Growth/size analysis of N2 and N2^*peel-1(null)*^. The body size of young adult animals were measured at 72 hours and normalized to the average size of N2. **E. - F.** Competition assays between indicated strains in standard laboratory conditions; positive values indicate Strain 1 is more fit and negative values indicate Strain 2 is more fit. **E.** Competition between the wild-type N2 *peel-1* allele and the *peel-1* loss-of-function mutation indicate a fitness benefit for *peel-1* (in assays with the marker in both backgrounds), which accounts for 32% of the difference arising from the relative fitness of the CB4856 introgression of *zeel-1;peel-1*. The relative fitness of N2^*npr-1;glb-5(CB4856)*^ over N2^*marker*^ is shown as a positive control. **F.** *zeel-1* shows no effect on the fitness benefit conferred by *peel-1:* there was no difference in fitness between *peel-1* loss-of-function strains with and without the *zeel-1* loss-of-function mutation, and there was no difference in the relative fitness increase conferred by *peel-1* with or without the zeel-1 loss-of-function mutation. For all panels, box plot show quartiles of the dataset while the whiskers cover the entire distribution of the data minus outliers; ***p<0.001 and *p<0.05 by two-tailed t-test.

In the N2 background, the *peel-1* toxin is expressed in the sperm and delivered to the embryo, but suppressed by the presence of the *zeel-1* antidote expressed by the embryo (Seidel et al., 2011). To test whether the fitness advantage of *peel-1* is *zeel-1* dependent, we generated a null *zeel-1* allele that produces a truncated protein sequence of five amino acids via an early stop (**Figure 3A**). After crossing the double mutant to N2, ~25 % of selfed cross-progeny of the N2 / N2^*zeel-1(null);peel-1(null)*^ hybrids died, confirming antidote loss-of-function (**Figure 3B**). Competition experiments between N2^*zeel-1(null);peel-1(null)*^ and N2^*peel-1(null)*^ showed no fitness differences between them (**Figure 3F**), suggesting that *peel-1* increases fitness in a *zeel-1* independent pathway.

This is not necessarily surprising, as the role of *peel-1* in a secondary biological process was considered in its initial characterization (Seidel et al., 2011). Such a role would help the initial spread of the element during its formation, when its low frequency (where gene drive is ineffective) and its initial toxicity (before *zeel-1* could evolve to counteract it) should prevent its spread. Our work supports that model, suggesting that both roles of *peel-1* could co-evolve together. But then, why hasn’t the element fixed? The *peel-1;zeel-1* locus shows a signature of balancing selection, which appears widespread in *C. elegans.* Hyperdivergent regions, including that spanning *peel-1;zeel-1*, punctuate the genome; balancing selection across diverse ecological niches may explain their maintenance (Lee et al., 2021). Previously, maintenance of the *peel-1/zeel-1* element was hypothesized to arise from tight linkage to a nearby polymorphism under balancing selection (Seidel et al., 2011). Our results suggest that *peel-1* could be under balancing selection itself. *peel-1* confers a fitness benefit within the lab environment, and it may pleiotropically influence other life history traits or affect fecundity and growth rate differently in different environments, providing alternate fitness strategies for local adaptation.

Previous work has suggested that TA elements may shape evolution by promoting selfing, to escape the cost of selfish gene drive (Noble et al., 2021). Here we provide a mechanism for their spread and maintenance that helps to explain their prevalence in selfing *Caenorhabditis* (Ben-David et al., 2021; Noble et al., 2021; Sweigart et al., 2019). Moreover, our observation of a toxin directly affecting biological traits mirrors work in transposable elements, which are also selfish elements that can be domesticated for phenotypic benefit to the organism (Werren, 2011). We posit that these findings demonstrate an outsized role for TA elements in shaping evolutionary trajectories.

## CONCLUSION

We have brought genomic editing and experimental evolution resources to bear on the study of a toxin-antidote element, addressing long-standing questions about their origin and maintenance in populations. We discovered that *peel-1* plays a biological role outside of its role as a toxin, affecting growth, fecundity, and fitness of non-hybrid genotypes, supporting recent arguments that non-selfish activity in inbred lineages may explain the prevalence of TA elements in non-obligate outcrossers (Noble et al., 2021; Sweigart et al., 2019). To our knowledge, this is the first measurement of the fitness cost of a TA element to the host and the first demonstration that a TA element can benefit the organism. We believe that other TA elements identified in *Caenorhabditis* species will also play biological roles, explaining how they can be retained in non-outcrossing populations.

## METHODS

### Growth conditions

Strains were cultivated on agar plates seeded with *E. coli* strain OP50 at 20°C (Brenner, 1974). The following strains were used in the study:

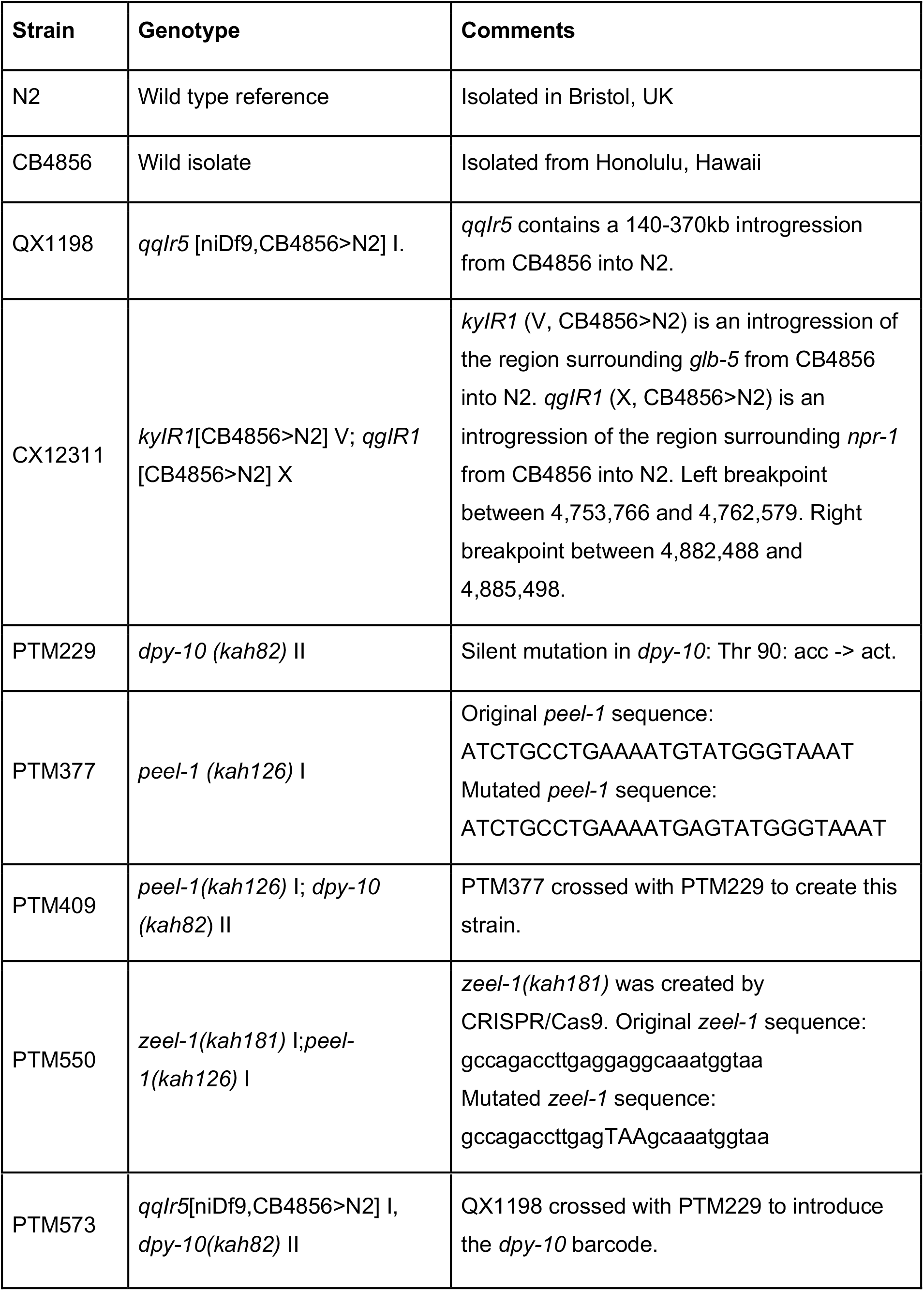

CRISPR/Cas9 was used following a previously published co-conversion method to edit the target gene and *dpy-10* gene at the same time (Arribere et al., 2014). The following primers/sequences were used to create the CRISPR/Cas9 strains:

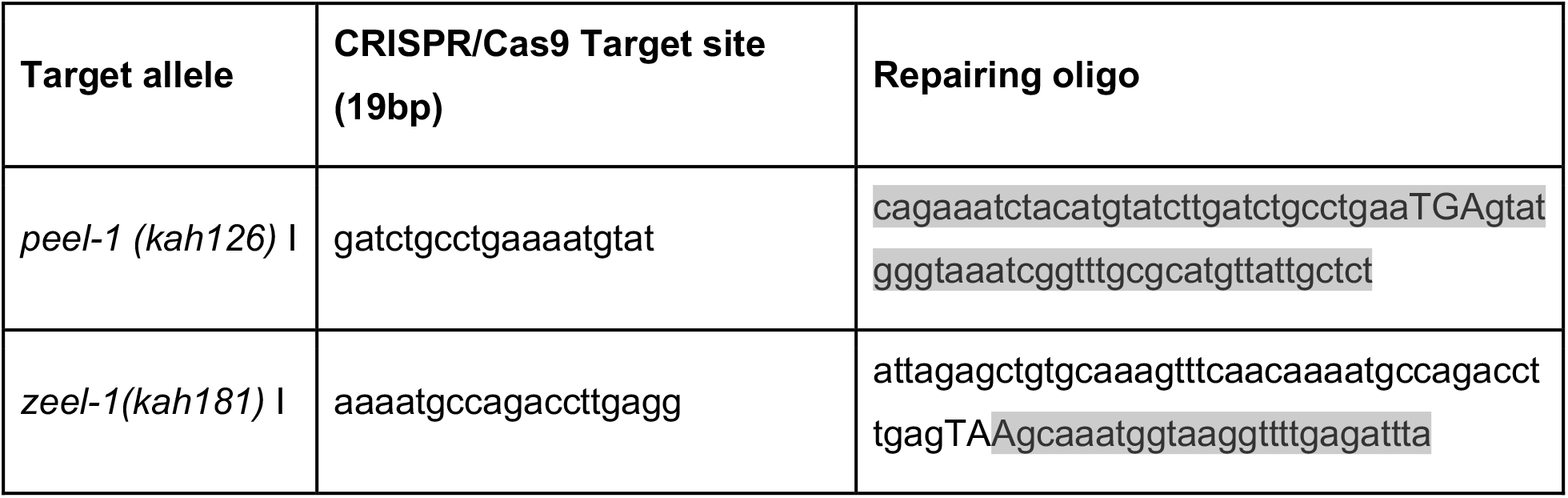

### Population dynamics prediction

All code to control population dynamics parameters and then plot the trajectories were stored at https://github.com/lijiang-long/TA_modeling. To calculate the allele frequency change at different frequencies of *peel-1/zeel-1*, the population is initiated with Hardy Weinberg equilibrium such that the frequency of homozygous *peel-1/zeel-1* is the square of its allele frequency, and so on and so forth. The frequency of each genotype is updated each generation using the family-based toxin-antidote evolution dynamics in Table S1. This population is allowed to evolve 5 generations to deviate from Hardy Weinberg equilibrium and reach the evolution trajectory of *peel-1/zeel-1.* The population evolves another generation, and the allele frequency change in this generation is used for plotting. To generate the heatmap where the frequency of *peel-1/zeel-1* after 1k generations is plotted against varying outcrossing rate and fitness cost, the population is initiated with half *peel-1/zeel-1* allele. The genotype frequency is calculated assuming Hardy Weinberg equilibrium. The population then evolves 1000 generations following Table S1. The final allele frequency of *peel-1/zeel-1* is then plotted on the heatmap.

### Competition assay to measure organism fitness

Competition experiments followed previous work (Zhao et al., 2018). All pairwise competition assays were performed on 9 cm NGM plates, seeded with OP50 bacteria and stored at 4°C until 24 hours before use. At the beginning of the experiment, 10 L4 worms of each strain were transferred onto the same plate. This plate was then incubated at 20°C for 5 days. To propagate the next generation, a 1 cm agar chunk was transferred to a new 9 cm NGM plate. The old plate was then washed with 1 ml of M9 buffer to collect worms and stored at −80°C. Subsequently, this transfer and collection procedure was held every three days for a total of 7 transfers. The genomic DNA from the 1^st^, 3^rd^, 5^th^ and 7^th^ transfer was isolated using Zymo 96-well DNA isolation kit (cat #D4071). Isolated genomic DNA was fragmented using EcoRI-HF by incubation at 37°C for 4 hours and purified using a Zymo 96-well DNA purification kit (cat #D4024). After purification, DNA concentrations were measured using Qubit DNA HS assay and adjusted to 1ng/μL. To quantify the relative proportion of the two strains, a previously designed Taqman probe was used targeting the *dpy-10* gene. After this, the DNA and Taqman probe were mixed with the digital plate PCR (ddPCR) mix and processed through standard ddPCR procedures. The fractions of each strain were quantified using the BioRad QX200 machine with standard absolute quantification protocol. To estimate relative fitness, a linear regression model was applied to the DNA proportion data using the following equation with the assumption of one generation per transfer.

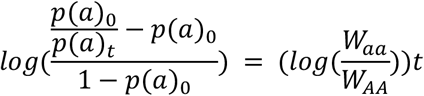

### Fecundity assays

Fecundity assays were performed at 20°C using 3 cm NGM plate seeded with 50 μL of OP50 bacteria with OD_600_ of 2.0. The plates were allowed to dry overnight and stored at 4°C until 24 hours before use. At the beginning of the assay, six fourth larval stage (L4) worms were transferred to each assay plate. The worms were allowed to grow and lay eggs for the first 24 hours after the assay began before being transferred to a new plate. This process was repeated every 12 hours thereafter until animals ceased laying eggs. The number of eggs laid was counted using a standard dissecting microscope. This process is repeated every 12 hours thereafter until 100 hours or there is no egg on the new plate. The average fecundity was calculated by summing over all time points and dividing by the total number of worms in a single assay plate.

### Growth rate assay

Growth rate assays were performed on standard NGM plates seeded with OP50 bacteria as previously described (Large et al., 2016). At the beginning of the assay, 10-20 adult worms were transferred onto an assay plate to lay eggs. After 2 hours, they were transferred off of the plate, leaving ~80 eggs per plate. The plates were incubated for 72 hours at 20°C. At this point, the assay plate was mounted onto a video tracking camera and recorded for one minute. The video clip was analyzed using a customized MATLAB script that tracks each animal and calculates the average size of each worm. The average size from each plate was then normalized by the average size of three N2 plates.

### Statistics

Significant differences between means were determined using unpaired, two-tailed t-tests assuming equal variance.

## ACKNOWLEDGEMENTS

We wish to acknowledge the core facilities at the Parker H. Petit Institute for Bioengineering and Bioscience at the Georgia Institute of Technology for the use of their shared equipment, services and expertise. Some strains were provided by the CGC, which is funded by NIH Office of Research Infrastructure Programs (P40 OD010440). We also thank the Kruglyak lab (UCLA) for strains. This research was supported in part through research cyberinfrastructure resources and services provided by the Partnership for an Advanced Computing Environment (PACE) at the Georgia Institute of Technology. This research was funded by NIH grant R35 GM119744 to A.B.P and NIH grant R35 GM139594 to P.T.M.

## Figure S1

**Figure S1.**
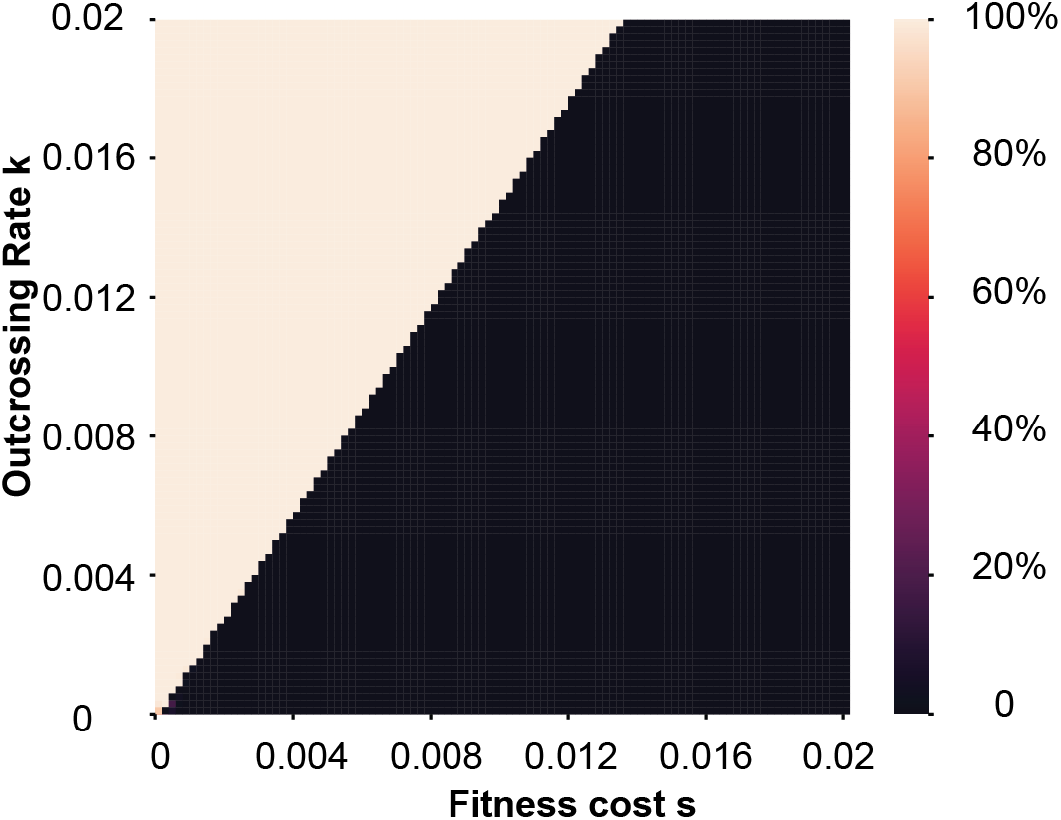
Heat map of *peel-1/zeel-1* frequency after 100 generations. The x-axis shows carrying costs and the y-axis shows outcrossing rates over a range typical of *C. elegans* in nature. Initial frequency was 50%.

**Table S1.** A family-based model for the *peel-zeel* evolution dynamics.

| Family | Mating types |  | Frequency | Female fitness | Offspring genotype |  |  |
| --- | --- | --- | --- | --- | --- | --- | --- |
|  | Sire | Dam |  |  | PP | P+ | ++ |
| 1 | PP | PP | $X_{pp}X_{pp}k$ | 1-s | 1 | | |
| 2 | P+ | PP | $X_{p+}X_{pp}k$ | 1-s | 0.5 | 0.5 | |
| 3 | ++ | PP | $X_{++}X_{pp}k$ | 1-s | | 1 | |
| 4 | PP | P+ | $X_{pp}X_{p+}k$ | 1-hs | 0.5 | 0.5 | |
| 5 | P+ | P+ | $X_{p+}X_{p+}k$ | 1-hs | 0.25 | 0.5 | $0.25(1-t)$ |
| 6 | ++ | P+ | $X_{++}X_{p+}k$ | 1-hs | | 0.5 | 0.5 |
| 7 | PP | ++ | $X_{pp}X_{++}k$ | 1 | | 1 | |
| 8 | P+ | ++ | $X_{p+}X_{++}k$ | 1 | | 0.5 | $0.5(1-t)$ |
| 9 | ++ | ++ | $X_{++}X_{++}k$ | 1 | | | 1 |
| 10 | PP selfing | | $X_{pp}(1-k)$ | 1-s | 1 | | |
| 11 | P+ selfing | | $X_{p+}(1-k)$ | 1-hs | 0.25 | 0.5 | $0.25(1-t)$ |
| 12 | ++ selfing | | $X_{++}(1-k)$ | 1 | | | 1 |

Parameter X denotes the ratio of a certain genotype in a population. Genotype P denotes *peel/zeel* and + denotes 'no *peel/zeel*'. The parameter k specifies the outcrossing rate. When k=1, there is complete outcrossing, and partial outcrossing is given by $0 < k < 1$ . The parameter s is the degree *peel-1/zeel-1* might reduce female fecundity. Dominance of the fecundity loss is defined by h. The parameter t models the paternal effect lethality. In the *peel-1/zeel-1* case, t is very close to 1.

